# A Large-Scale Foundation Model for RNA Function and Structure Prediction

**DOI:** 10.1101/2024.11.28.625345

**Authors:** Shuxian Zou, Tianhua Tao, Sazan Mahbub, Caleb N. Ellington, Robin Algayres, Dian Li, Yonghao Zhuang, Hongyi Wang, Le Song, Eric P. Xing

**Author notes:** Work done during internship at GenBio AI.

## Abstract

Originally marginalized as an intermediate in the information flow from DNA to protein, RNA has become the star of modern biology, holding the key to precision therapeutics, genetic engineering, evolutionary origins, and our understanding of fundamental cellular processes. Yet RNA is as mysterious as it is prolific, serving as an information store, a messenger, and a catalyst, spanning many undercharacterized functional and structural classes. Deciphering the language of RNA is important not only for a mechanistic understanding of its biological functions but also for accelerating drug design. Toward this goal, we introduce AIDO.RNA, a pre-trained module for RNA in an AI-driven Digital Organism [1]. AIDO.RNA contains a scale of 1.6 billion parameters, trained on 42 million non-coding RNA (ncRNA) sequences at single-nucleotide resolution, and it achieves state-of-the-art performance on a comprehensive set of tasks, including structure prediction, genetic regulation, molecular function across species, and RNA sequence design. AIDO.RNA after domain adaptation learns to model essential parts of protein translation that protein language models, which have received widespread attention in recent years, do not. More broadly, AIDO.RNA hints at the generality of biological sequence modeling and the ability to leverage the central dogma to improve many biomolecular representations. Models and code are available through ModelGenerator in https://github.com/genbio-ai/AIDO and on Hugging Face.

## 1 Introduction

RNA, an essential biomolecule found in all living organisms, holds the distinction of being considered the original molecule of life [2]. Its significance extends beyond its role in bridging genetic information and protein synthesis, as it plays a crucial part in various cellular processes such as metabolism, transport, signaling, and regulation. Messenger RNA (mRNA) carries genetic instructions, transfer RNA (tRNA) aids in translating mRNA into amino acids, and ribosomal RNA (rRNA) forms an integral part of ribosomes involved in protein synthesis. Furthermore, small non-coding RNA molecules, such as microRNA and small interfering RNA, regulate gene expression by silencing or degrading specific mRNA molecules. Understanding the emergence of diverse structures and functions from a simple 4-letter chemical vocabulary is vital for comprehending cellular processes, genetic regulation, and disease mechanisms. RNA’s synthesizability, programmability, and broad functionality make it an attractive candidate for therapeutic development and metabolic engineering [3].

However, due to the dynamic nature of RNA structures, there are only a few thousand RNA structure data available in the Protein Data Bank [4], making it difficult for RNA to get its AlphaFold [5] moment [6]. Furthermore, functional labels specific to RNA tasks are often scarce. Despite the scarcity of RNA structural and functional data, the rapid progress in next-generation sequencing technology has led to a substantial accumulation of RNA sequence data. Similar scenarios have been observed in the fields of Natural Language Processing (NLP) and protein science, where substantial quantities of unannotated sequences are accessible. Inspired by the huge success of foundation models in the two domains, we seek a foundation model for RNA, aiming to benefit a diverse set of RNA-related tasks, including RNA structure/function prediction and RNA sequence design.

In recent years, several RNA FMs have been proposed [11, 12, 13, 9, 14, 15, 16, 17, 7, 10, 8, 18], most of which are encoder-only transformers pre-trained using the Masked Language Modeling (MLM) objective [19, 20] (See Appendix E). These models have shown impressive results in RNA secondary structure prediction and function prediction [9, 10], demonstrating the potential of large language models (LLMs) in the RNA domain. However, existing RNA foundation models are relatively small (up to 650M parameters) compared to protein language models (up to 100B parameters) [10, 21]. Scaling LLMs for RNA remains an interesting and open challenge.

Furthermore, translating sequence modeling methods from NLP to RNA requires substantial improvements beyond protein language models. While LLM technology is directly applicable to RNA, determining the ideal pre-training dataset for general-purpose RNA foundation models remains unresolved. Unlike the protein domain, where UniRef [22] is a typical pre-training data source, RNA sequences are scattered across various biological databases. Currently, there exist two easily accessible RNA sequence databases: 1) RNAcentral, a high-quality ncRNA sequence database with 42 million samples, and 2) MARS, a noisy RNA sequence database containing 1.7 billion sequences. We investigate both datasets for pre-training purposes and discover that utilizing a high-quality database is crucial, resulting in improved overall downstream performance (See Section 3.5).

We introduce AIDO.RNA, a 1.6B-parameter RNA foundation model pre-trained on 42 million ncRNA sequences from RNAcentral. To the best of our knowledge, AIDO.RNA is the largest RNA foundation model to date. To evaluate its performance, we create a comprehensive RNA sequence understanding benchmark comprising 26 datasets from 9 task categories, including structure, function, and mRNA-related tasks relevant to mRNA vaccine design. As shown in Figure 1, AIDO.RNA surpasses previous state-of-the-art (SOTA) results on 24 out of 26 tasks. In particular, our model excels in RNA secondary structure prediction and translation efficiency prediction, which are tasks specifically relevant to the RNA level, as opposed to DNA or protein levels. Furthermore, we evaluate AIDO.RNA on the 3D RNA inverse design, which involves generating sequences based on 3D RNA backbones. Experiment results show that our model enhances performance compared to the previous SOTA method, gRNAde [23], with or without fine-tuning. These results demonstrate AIDO.RNA’s strong capabilities in RNA language understanding and generation, positioning it as a potent foundation model for diverse RNA tasks.

**Figure 1.**
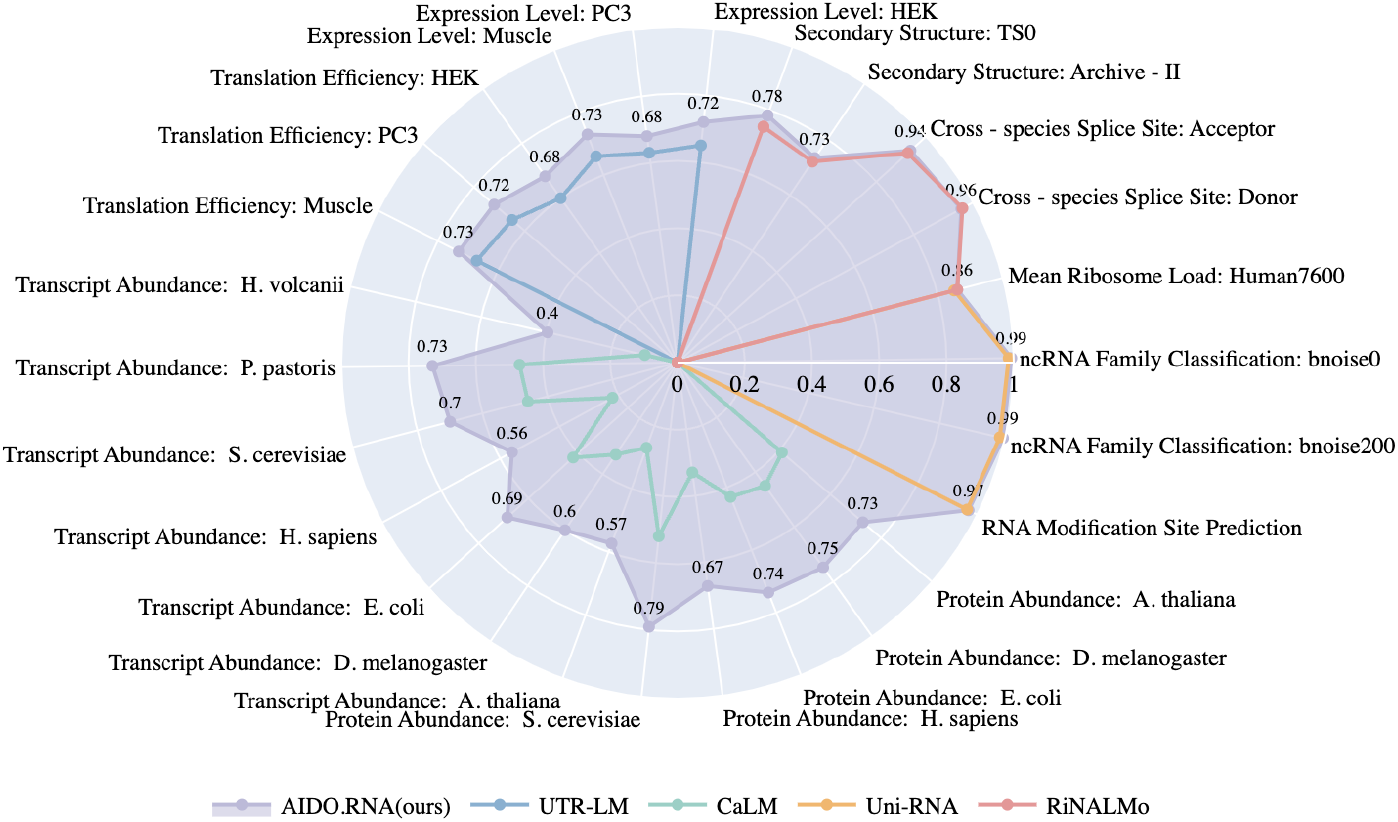
AIDO.RNA achieves SOTA results on 24 out of 26 RNA sequence understanding tasks. We compare our model with two domain expert models UTR-LM [7] and CaLM [8], and two general models Uni-RNA [9] and RiNALMo [10]. For all tasks, higher metric values indicate better performance.

### 2 Pre-training AIDO.RNA

In this work, we explore scaling up RNA foundation models. We adopt encoder-only transformer as our model architecture and use masked language modeling (MLM) as the pre-training objective. Special focus is given to the pre-training data, which remains under-explored and lacks consensus in the RNA domain. Our exploration in the pre-training data suggests that data quality outweigh data quantity, as illustrated in Section 3.5. Therefore, we leverage high-quality RNA sequences from RNAcentral [24] as the pre-training data and pre-train the first RNA foundation model at the scale of 1.6 billion parameters.

#### Pre-training data

RNAcentral database contains a comprehensive non-coding RNA sequence collection representing all ncRNA types from a broad range of organisms. In specific, we collect sequences from `rnacentral_active.fasta.gz and rnacentral_inactive.fasta.gz` from version 24.0 and then remove duplications using `SeqKit` toolkit. The resulting dataset contains 42 million unique ncRNA sequences. Notably, we do not use clustered sequences as RiNALMo [10] did for two reasons: 1) The similarity between sequences in a carefully curated dataset represents evolutionary selection and conservation, which should be kept for the model to learn; 2) It is very likely that clustering won’t help since the average cluster size is just 2.2 when we cluster the sequences to 0.7 sequence identity using MMseqs2 [25]. Furthermore, we find that the data follows a long tail distribution in terms of RNA types, as shown in Appendix Table 12. To have a better understanding of the generalization ability of our model on different RNA types, we downsample the frequent types and upsample the infrequent types for validation and testing. Distributions of the train, validation, and test set are shown in Appendix Table 12. The training dataset consists of 41.5M distinct ncRNA sequences, comprising a total of 30 billion nucleotides. On average, each sequence has 728 nucleotides.

#### Sequence tokenization

We encode each nucleotide (`A, T, C, G`) as a token and use `N` to represent other rare bases (`U` has been transformed to `T` in our dataset). We also introduce some special tokens, including [CLS], [SEP], `[MASK],` `[PAD]`. The vocabulary size is set to 16. When processing each RNA sequence, we prepend the `[CLS]` token at the beginning and append the `[SEP]` token at the end. This allows the model to separate a full-sized RNA from a cropped one.

#### Model architecture

Following common practice in the literature as summarized in Appendix Table 22, we adopt encoder-only transformer as our pre-training model architecture to extract meaningful biological representations from RNA sequences [19, 20]. Figure 2 illustrates our pre-training model architecture. In experiment, our 1.6B model AIDO.RNA contains 32 layers and 32 attention heads. The hidden size is set to 2,048 and the feed-forward hidden size is 5,440. We use Rotary Position Embedding (RoPE) [26] to allow better position modeling. In addition, we use LayerNorm [27] and SwiGLU activation function in our model to make it more expressive and stable in pre-training.

**Figure 2.**
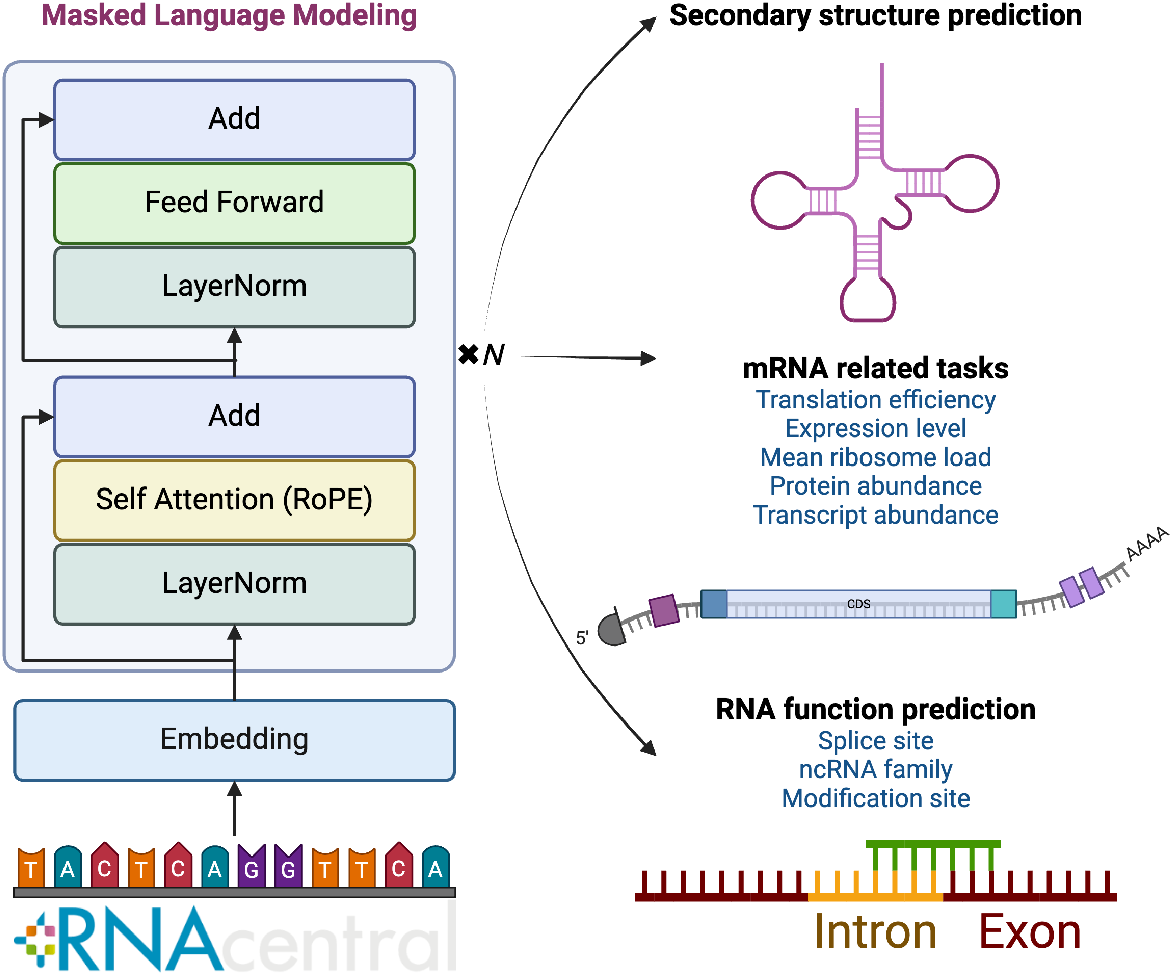
Pre-training model architecture of AIDO.RNA. AIDO.RNA takes masked sequences from RNAcentral as input and aims to reconstruct the masked tokens through MLM objective. After pre-training, the model can be applied to various downstream tasks. Figure created in BioRender.com.

#### Pre-training setting

We pre-train two models with different sizes, i.e., 650M, 1.6B, on non-coding RNA sequences. Unless otherwise specified, AIDO.RNA denotes the 1.6B one. We use the MLM objective with a masking ratio 0.15. In specific, this involves randomly selecting 15% of the input tokens for each input sequence. For the selected tokens, there are three possible operations: 1) The token is masked with a probability of 0.8; 2) The token is replaced with a random token with a probability of 0.1; 3) The token remains unchanged with a probability of 0.1. Cross entropy loss is computed on those selected tokens. We train our models on 30 billion unique nucleotides for 6 epochs. We use AdamW optimizer with weight decay of 0.01 [28]. The peak learning rate is set to 5e-5 and gradually decay to 1e-5 via cosine learning rate scheduler. All hyperparameters for pre-training are summarized in Appendix Table 13. We implement our code using the Megatron-LM framework. To accelerate pre-training, we use FlashAttention-2 [29] and use BFloat16 mixed precision training.

## 3 Results

### 3.1 AIDO.RNA captures RNA structural information

As with proteins, structure determines RNA function. RNA secondary structure, formed by base pairing, is more stable and accessible than its tertiary form within cells. Accurate prediction of RNA secondary structure is essential for tasks such as higher-order structure prediction and function prediction [30]. We utilize two benchmark datasets created by Singh et al. (2019) [31] and Szikszai et al. (2022) [32] for RNA secondary structure prediction. The first dataset, derived from bpRNA- 1m [33], is divided into three splits: TR0 for training, VL0 for validation, and TS0 for testing. The second dataset, which is used for generalization assessment, contains nine distinct RNA families. Following RiNALMo, we use the same metric calculation approach proposed by [34]. We consider (*i* ± 1, *j*) and (*i, j* ± 1) pairings as correct predictions for a nucleotide pairing (*i, j*), where *i* and *j* denote nucleotide index in the RNA sequence.

Table 1 shows the results of our model and several baselines on the bpRNA-TS0 test set. AIDO.RNA achieves SOTA results on this dataset, with an F1 score of 0.787, outperforming RNAErnie [18] and RiNALMo by large margins. This result indicates that our model pre-trained on sequence data learns substantially more structural information than previous methods. Furthermore, a case study detailed in Appendix Section D reveals that our model learns functional dependencies within RNA sequences without labeled data, as shown in Figure 3.

**Table 1:** RNA secondary structure prediction results on bpRNA-TS0.

|  | Precision | Recall | F1-score |
| --- | --- | --- | --- |
| SPOT-RNA [35] | 0.594 | 0.693 | 0.619 |
| UFold [36] | 0.607 | 0.741 | 0.654 |
| RNA-FM [12] | 0.709 | 0.664 | 0.676 |
| RNAErnie[18] | 0.575 | 0.678 | 0.622 |
| RiNALMo [10] | 0.784 | 0.730 | 0.747 |
| AIDO.RNA(ours) | **0.815** | **0.769** | **0.783** |

**Figure 3.**
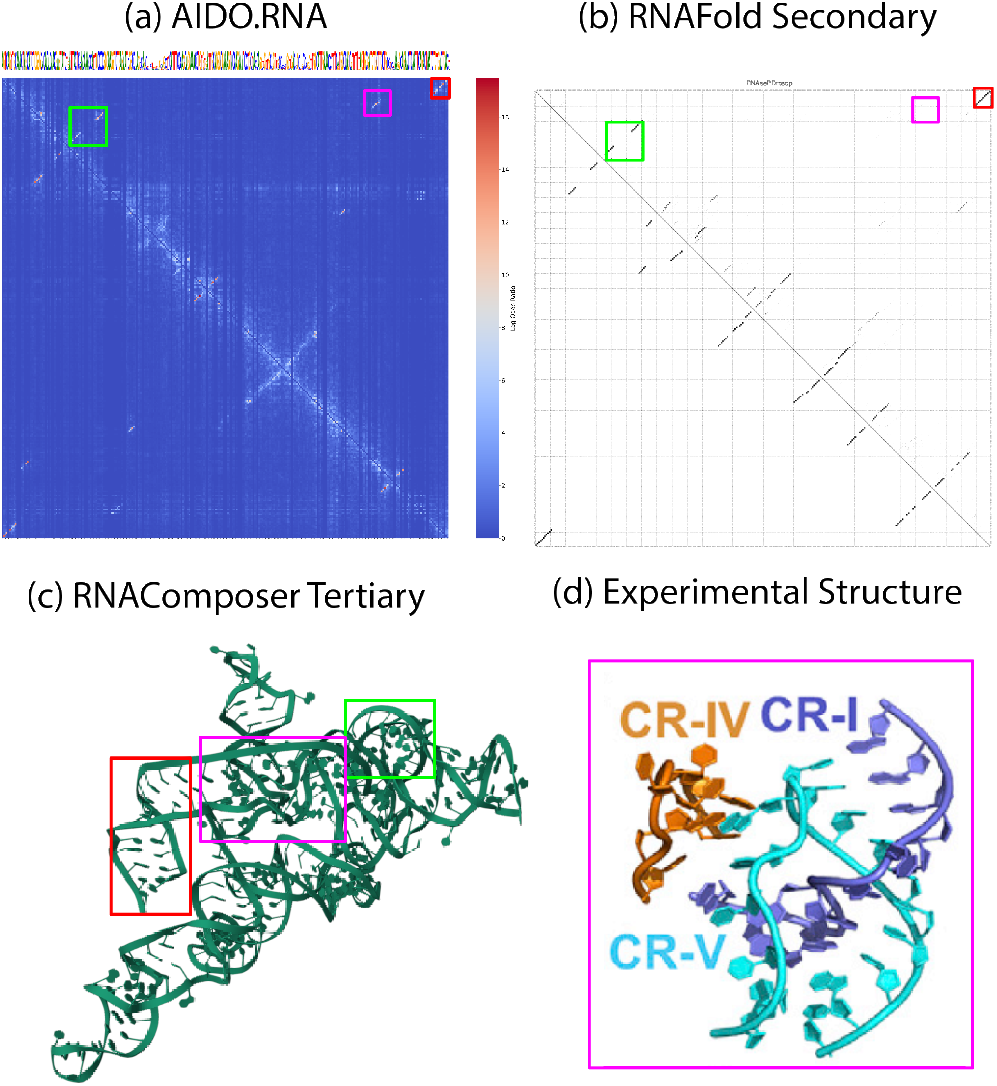
Unsupervised pre-training captures secondary and tertiary structures in the RNAseP component of the eukaryotic ribosome. Left: predictions; Right: ground truths.

We further test the generalization ability of AIDO.RNA using the dataset from [32]. We use 9-fold cross-validation, with each fold corresponding to one RNA family. We compare our model with RNA-FM [12], RNAstructure [37], CONTRAfold [38], UFold [36], and MXfold2 [39]. These models are trained and tested on the split datasets, with CONTRAfold using EternaFold parameters from the EternaBench dataset. Table 2 shows the inter-family generalization results in RNA secondary structure prediction. Our model outperforms RNAstructure in 6 out of 9 families and achieves the second-highest scores in two families.

**Table 2:**
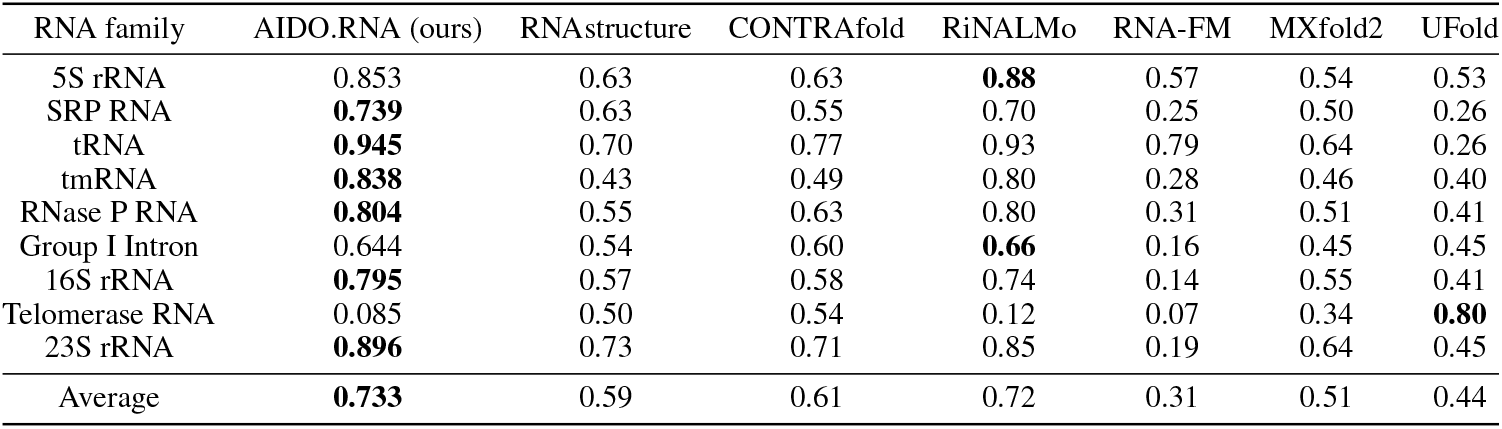
Inter-family generalization for secondary structure prediction on filtered Archive-II. Reported is the average F1 score. Bold denotes the best performance within a family.

### 3.2 AIDO.RNA facilitates genetic engineering

Genes introduced to a cell via genetic engineering often must be expressed as proteins to affect cellular functions. mRNA plays a vital role in protein synthesis by transferring genetic information from DNA to ribosomes for protein production. It consists of three main regions: the 5’ untranslated region (5’ UTR), coding sequence (CDS), and 3’ untranslated region (3’ UTR), each serving specific functions in mRNA regulation. Understanding their effect on transcription and translation is essential for improving the success of genetic engineering and gene therapies.

#### 3.2.1 mRNA translation efficiency and expression level prediction based on 5’ UTR

Protein expression is highly dependent on 1) the relative abundance of the mRNA transcript in the cell (refer as mRNA expression level), and 2) the rate at which mRNA molecules are translated into proteins within a cell (refer as mRNA translation efficiency). Following UTR-LM [7], we use the same datasets and metrics for mRNA translation efficiency and expression level prediction. Specifically, mRNA expression level is quantified using RNA-sequencing RPKM (reads per kilobase of transcript per million mapped reads), while mRNA translation efficiency is determined by dividing Ribo-seq RPKM by RNA-sequencing RPKM. To evaluate these tasks, we employ three datasets collected from human muscle tissue (muscle), the human prostate cancer cell line PC3 (PC3), and the human embryonic kidney 293T cell line (HEK).

We fully fine-tune our model on each of the cell line dataset using 10-fold cross-validation. We adopt Spearman correlation coefficient as the evaluation metric. Table 3 shows the results of our model and baseline models, including the domain expert UTR-LM which is pre-trained on 5’ UTR sequences. On the mRNA translation efficiency prediction task, AIDO.RNA achieves SOTA on each cell line, outperforming UTR-LM by large margins. On average across the three cell lines, AIDO.RNA attains a Spearman correlation coefficient of 0.71, with a relative improvement over UTR-LM of +10.9%. Similarly, our model achieves SOTA results on all cell lines in the mRNA expression level prediction task. On the one hand, these results showcase the superior performance of our model on the two types of tasks. On the other hand, they indicate that our model has strong generalization capabilities, successfully adapting to a new domain beyond its original pre-training scope.

**Table 3:** mRNA translation efficiency and expression level prediction results. We use 10-fold cross-validation. Reported is the average Spearman correlation coefficient across 10 folds.

|  | mRNA translation efficiency |  |  |  | mRNA expression level |  |  |  |
| --- | --- | --- | --- | --- | --- | --- | --- | --- |
|  | Muscle | PC3 | HEK | AVG | Muscle | PC3 | HEK | AVG |
| Optimus 5-Prime [40] | 0.41 | 0.38 | 0.36 | 0.38 | 0.15 | 0.19 | 0.18 | 0.17 |
| Cao-RF [41] | 0.63 | 0.63 | 0.55 | 0.60 | 0.64 | 0.59 | 0.57 | 0.60 |
| RNABERT [11] | 0.56 | 0.48 | 0.41 | 0.48 | 0.60 | 0.45 | 0.47 | 0.51 |
| RNA-FM [12] | 0.30 | 0.32 | 0.23 | 0.28 | 0.30 | 0.20 | 0.22 | 0.24 |
| UTR-LM [7] | 0.67 | 0.65 | 0.60 | 0.64 | 0.66 | 0.63 | 0.65 | 0.65 |
| AIDO.RNA (ours) | **0.73** | **0.72** | **0.68** | **0.71** | **0.73** | **0.68** | **0.72** | **0.71** |

#### 3.2.2 Mean ribosome load prediction based on 5’ UTR

Ribosomes are cellular structures responsible for protein synthesis, and the ribosome load on an mRNA molecule can influence the rate and efficiency of protein production, and the success of genetic engineering. Predicting ribosome load can provide valuable insights into gene expression regulation, translation efficiency, and cellular processes. Due to its significance, several studies emphasize on computational prediction of mean ribosome load (MRL), which is defined as the number of ribosomes bound to a specific mRNA molecule at any given time [40, 12, 9]. We use datasets from [40], which include 5’ UTR sequences with measured MRL values. The validation and test sets, namely Random7600 and Human7600, are generated by sampling sequences of varying lengths (25-100 nucleotides) from random and human UTR sequences. Each length category contains 100 sequences with the deepest read coverage. The remaining random 5’ UTRs with sufficient read coverage formed the training dataset. As shown in Table 4, our model fine-tuned on the dataset attains the same *R*^2^ score as the SOTA model RiNALMo.

**Table 4:** Mean ribosome load prediction results.

| | $R^2$ score ↑ |
| --- | --- |
| Optimus 5-Prime [40] | 0.78 |
| RNA-FM [12] | 0.79 |
| Uni-RNA [9] | 0.85 |
| RiNALMo [10] | **0.86** |
| AIDO.RNA (ours) | **0.86** |

#### 3.2.3 Transcript abundance prediction based on CDS

Transcript abundance refers to the quantity of a specific RNA transcript within a cell or tissue at a given time. It represents the amount of mRNA molecules produced from a particular gene and serves as an indicator of gene expression. We leverage the transcript abundance datasets from CaLM [8], which contain samples from seven organisms. Notably, although sharing conceptual similarities with the mRNA expression level prediction task described in Section 3.2.1, this task diverges in terms of input requirements. Instead of utilizing 5’ UTR sequences, it focuses on coding sequences. In essence, this task aligns more closely with protein-level tasks rather than RNA-level tasks.

We fine-tune our models on each of the seven organisms using LoRA [42]. We use 5-fold cross-validation and adopt the Pearson correlation coefficient as the evaluation metric, following the setting in CaLM. Table 5 shows the results of our models and the baselines. AIDO.RNA outperforms the codon language model CaLM by large margins on all organisms. On average across the seven organisms, it achieves a 0.560 Pearson correlation coefficient, outperforming CaLM by an absolute improvement of 0.236. When comparing to the protein language model ESM2-650M [43], our model also achieves better results on most of the organisms, indicating that the nucleotide space provides additional information compared to the amino acid space.

**Table 5:** Transcript abundance prediction results. We use 5-fold cross-validation for each dataset. Reported is the average Pearson correlation coefficient across 5 folds. AIDO.RNA-CDS denotes our CDS domain-adaptive model.

|  | *A. thaliana* | *D. melanogaster* | *E.coli* | *H. sapiens* | *S. cerevisiae* | *P. pastoris* | *H. volcanii* | AVG |
| --- | --- | --- | --- | --- | --- | --- | --- | --- |
| CaLM [8] | 0.270 | 0.330 | 0.420 | 0.220 | 0.460 | 0.470 | 0.100 | 0.324 |
| ESM2-650M(LoRA) | 0.460 | 0.488 | 0.430 | 0.449 | 0.672 | 0.602 | 0.269 | 0.482 |
| AIDO.RNA(ours) | 0.510 | 0.535 | 0.632 | 0.527 | 0.656 | 0.684 | 0.377 | 0.560 |
| AIDO.RNA-CDS(ours) | **0.573** | **0.601** | **0.685** | **0.560** | **0.699** | **0.729** | **0.397** | **0.606** |

To better adapt our model from the ncRNA domain to the CDS domain, we continue to pre-train our model on 9 million CDS sequences from CaLM [44]. Intriguingly, our domain-adaptive model achieves impressive performance gain across all datasets over the pre-trained model, setting new SOTAs on these tasks. These results suggest that: 1) Patterns learned from ncRNA sequences lay a solid foundation for generalization to the coding sequence region, and 2) Continued pre-training proves to be an effective strategy in the RNA domain.

#### 3.2.4 Protein abundance prediction based on CDS

Protein abundance refers to the quantity of a specific protein present within a cell or tissue at a given time. Analyzing protein abundance provides insights into protein expression patterns, cellular processes, and regulatory mechanisms. We utilize protein abundance datasets from CaLM, encompassing five organisms. The abundance labels are estimated as the number of protein copies per cell, as annotated in PAXdb [45]. In machine learning, this task is formulated as a sequence-level regression problem. We use the same fine-tuning and evaluation schemes for these tasks as in Section 3.2.3. As shown in Table 6, our model achieves SOTA performance on most datasets, in line with the results in transcript abundance prediction tasks. This consistency across tasks demonstrates that our model is competent in protein-level tasks, opening a new avenues for studying proteins.

**Table 6:** Protein abundance prediction results. We use 5-fold cross-validation for each dataset. Reported is the average Pearson correlation coefficient across 5 folds.

|  | *A. thaliana* | *D. melanogaster* | *E.coli* | *H. sapiens* | *S. cerevisiae* | AVG |
| --- | --- | --- | --- | --- | --- | --- |
| CaLM [8] | 0.410 | 0.450 | 0.430 | 0.330 | 0.520 | 0.428 |
| ESM2-650M(LoRA) | 0.689 | 0.662 | 0.595 | **0.682** | 0.682 | 0.662 |
| AIDO.RNA(ours) | 0.644 | 0.688 | 0.685 | 0.560 | 0.757 | 0.667 |
| AIDO.RNA-CDS(ours) | **0.728** | **0.748** | **0.736** | 0.671 | **0.791** | **0.735** |

### 3.3 AIDO.RNA predicts RNA function

#### 3.3.1 Cross-species splice site prediction

RNA splicing is a crucial step in gene expression, particularly in eukaryotic organisms. It is the process by which introns (non-coding regions) are removed from pre-messenger RNA (pre-mRNA) sequences, and the remaining exons (coding regions) are joined together to form mature mRNA. Predicting splice sites is essential for uncovering the structure of genes and gaining insights into the mechanisms of alternative splicing. Depending on the location in the pre-mRNA sequence, the splice site can be classified into two types: donor and acceptor. We leverage the dataset from Spliceator [46], which contains a donor dataset and an acceptor dataset. For each dataset, the task is formulated as a sequence-level binary classification task, which is to predict whether a given RNA sequence contains a donor/acceptor or not. We fine-tune AIDO.RNA on the donor and acceptor datasets separately using LoRA. We then test it on four unseen species that are not shown in the training data, including Zebrafish, fly, worm, and plant. Table 7 shows the average scores of the acceptor and donor dataset for our model and baselines. AIDO.RNA performs slightly better on 3 out of 4 species, with an average F1 score of 0.949 across four species.

**Table 7:** Cross-species splice site prediction results. The average F1 score across the donor and acceptor datasets is reported. Results of RiNALMo are reproduced by us by using their codebase and hyperparameters.

|  | Zebrafish | Fly | Worm | Plant | AVG |
| --- | --- | --- | --- | --- | --- |
| Spliceator [46] | 0.935 | 0.929 | 0.916 | 0.929 | 0.927 |
| DNABERT [47] | 0.951 | 0.931 | 0.909 | 0.909 | 0.925 |
| SpliceBERT [14] | 0.957 | 0.946 | 0.934 | 0.936 | 0.943 |
| Uni-RNA [9] | 0.964 | 0.950 | **0.939** | 0.936 | 0.947 |
| RiNALMo* [10] | **0.965** | 0.948 | 0.935 | 0.932 | 0.945 |
| AIDO.RNA (ours) | **0.965** | **0.949** | 0.936 | **0.945** | **0.949** |

#### 3.3.2 Non-coding RNA family classification

ncRNAs play important regulatory roles in various cellular processes. Depending on the sequence length, ncRNAs can be classified as short (≤ 200 nucleotides) or long (>200 nucleotides). Following [48], we leverage our model to predict short ncRNA functional families curated from Rfam [49] using only sequences as input. In machine learning, this task is a sequence-level classification task, with 88 classes in the label space. We assess the prediction performance under the uncertainty of where the ncRNA sequence starts and ends. We use the dataset from [48], which contains sequences with different levels of added boundary noise. In specific, a sequence with 0% boundary noise denotes the original ncRNA sequence. A sequence with 200% boundary noise refers to the addition of random nucleotides, equivalent to 100% of the sequence length, at both the beginning and the end of the ncRNA sequence. We fine-tune our model on datasets with different boundary noises using LoRA. Table 8 shows the results of our model and the baseline models. AIDO.RNA achieves a 0.993 accuracy on the dataset with 0% boundary noise, significantly outperforms Uni-RNA [9]. When dealing with 200% boundary noise, AIDO.RNA attains a SOTA score of 0.994, showing its robustness against boundary noise.

**Table 8:** ncRNA family classification results. Accuracy is reported.

|  | 0% boundary noise | 200% boundary noise | AVG |
| --- | --- | --- | --- |
| 1-mer-CNN [48] | 0.870 | 0.810 | 0.840 |
| 2-mer-CNN [48] | 0.880 | 0.840 | 0.860 |
| 3-mer-CNN [48] | 0.890 | 0.840 | 0.865 |
| Uni-RNA [9] | 0.985 | 0.984 | 0.985 |
| AIDO.RNA (ours) | **0.993** | **0.994** | **0.993** |

#### 3.3.3 RNA modification site prediction

Post-transcriptional RNA modifications are chemical modifications that occur on RNA molecules after transcription, which can alter the structure, stability, function, and processing of RNA molecules, playing crucial roles in various biological processes. Following [50], we assess our model’s ability to predict 12 types of RNA modification sites. In machine learning, this task is formulated as a sequence-level multi-label classification task. For evaluation, we compute the AUROC score for each modification site following the common practice in the literature. We fine-tune AIDO.RNA using LoRA. Table 9 shows the results of our model and the baseline models. AIDO.RNA achieves SOTA results on 12 types of modifications, with an average AUROC score of 0.971, outperforming MultiRM [50] and Uni-RNA [9].

**Table 9:** RNA modification site prediction results. AUROC score is reported

| | *Am* | *Cm* | *Gm* | *Tm* | *m*<sup>1</sup>*A* | *m*<sup>5</sup>*C* | *m*<sup>5</sup>*U* | *m*<sup>6</sup>*A* | *m*<sup>6</sup>*Am* | *m*<sup>7</sup>*G* | $\Phi$ | I | AVG |
| --- | --- | --- | --- | --- | --- | --- | --- | --- | --- | --- | --- | --- | --- |
| MultiRM [50] | 0.789 | 0.860 | 0.926 | 0.878 | 0.779 | 0.906 | 0.948 | 0.856 | 0.891 | 0.677 | 0.853 | 0.670 | 0.836 |
| Uni-RNA [9] | 0.929 | 0.968 | 0.986 | **0.959** | **0.954** | 0.976 | 0.958 | **0.994** | 0.978 | 0.956 | **0.942** | 0.993 | 0.966 |
| AIDO.RNA (ours) | **0.951** | **0.975** | **0.988** | 0.957 | **0.954** | **0.980** | **0.961** | **0.994** | **0.985** | **0.970** | 0.937 | **0.994** | **0.971** |

### 3.4 AIDO.RNA benefits 3D RNA inverse design

RNA sequence design involves the process of creating or generating RNA sequences with specific properties or functions [23, 51]. It is crucial to therapeutic innovation [52], synthetic biology [53, 54], and fundamental molecular biology research [55]. In this section, we extend AIDO.RNA with a discrete diffusion modeling framework to enable generative capabilities for RNA sequence design. We adopt a probabilistic diffusion approach [56, 57], iteratively refining sequences by masking and predicting optimal nucleotide compositions. It is a general method which supports both unconditional and conditional design. For details of our method, please refer to Appendix Section C.1.1.

We conduct experiments on RNA inverse folding, a task aiming to generate RNA sequences that fold into the given 3D structure [23, 58]. We use the dataset from Das et al. [59], which contains 4,025/100/98 train/validation/test samples. We evaluate our model in two settings: (1) adaptation with conditional diffusion where AIDO.RNA is fine-tuned for the inverse folding task; and (2) zero-shot generation where AIDO.RNA is frozen (refer to Appendix Section C for more details). As shown in Table 10, AIDO.RNA, integrated with the pipeline of a SOTA RNA inverse folding method, gRNAde [23], improves upon the original gRNAde in both zero-shot and diffusion-adaptation settings. In Appendix Table 21, we show results for 14 RNA structures of interest identified by Das *et al*. [59], where we can see that AIDO.RNA with diffusion-adaptation can enhance gRNAde’s performance by about 3%. We also provide a visualization of generated sequences for an example RNA (PDB ID: 3B58) in Appendix Figure 4, showing preserved structural details. These results showcase that AIDO.RNA can benefit RNA sequence design.

**Table 10:**
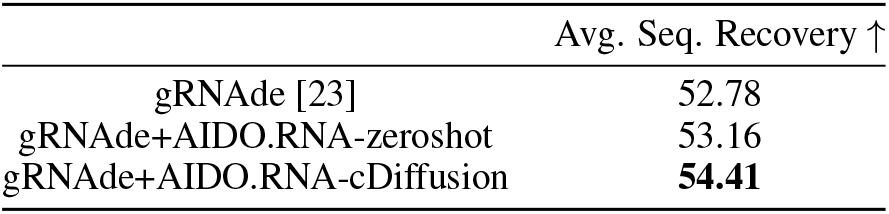
RNA inverse folding results.

**Figure 4.**
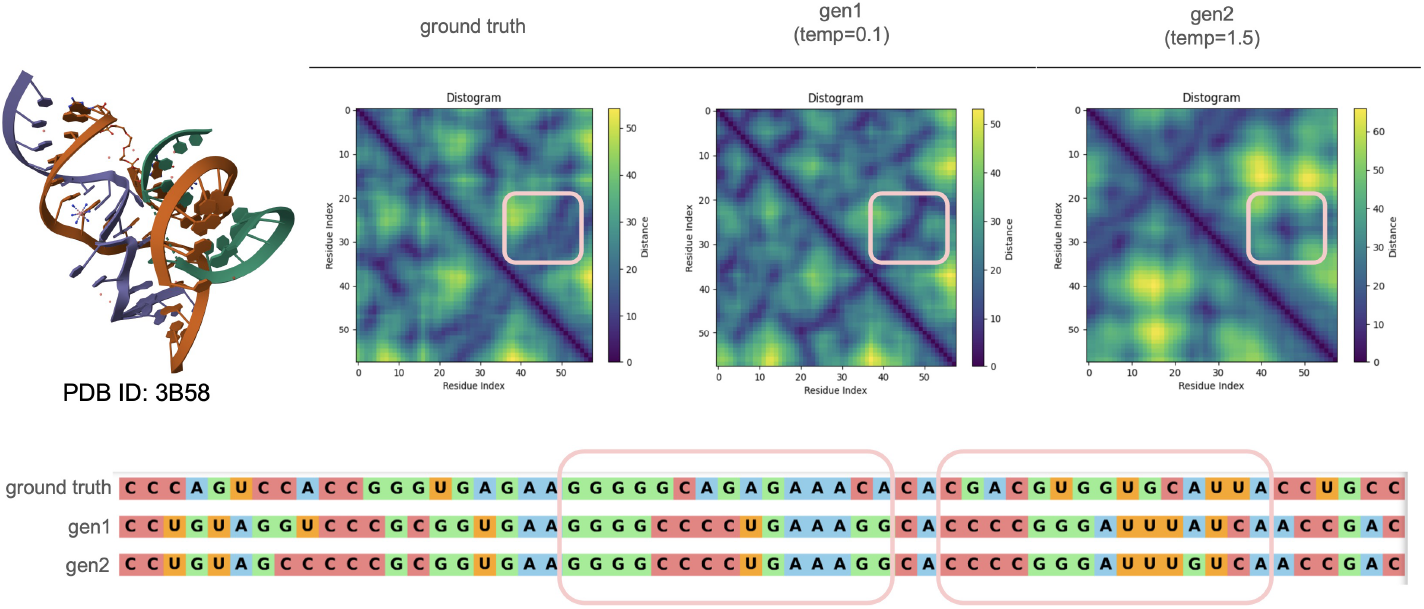
Generated sequence by our inverse folding framework leads to similar structures. Here we show the ground truth distogram (distances between base-pairs) of the given RNA 3D structure (PDB ID: 3B58) and the predicted distogram (by RhoFold) of two sequences generated by our framework. **Top row:** At the very left, we show the true structure of the RNA. The second image shows the ground truth distogram. The third image from the left (labeled as `gen1`) show the predicted distogram of RNA sequences generated by our framework, conditioned on the true structure, with low sampling temperature (temp=0.1). Note that sampling temperature controls how much randomness we allow for the generation (see [23] for details). We can see that, even though this distogram is a bit different from the ground truth, it shows structural similarity in several regions. We have marked a region of interest with rectangle for demonstration. The right-most image in the top row shows how the structure (represented as distrogram, denoted at `gen2`) changes when we allow more randomness in sampling, by introducing high sampling temperature (temp=1.5) [68, 69]. As expected, compared to the ground truth and `gen1`, the structural details are lost to a great extent in `gen2` as we allow more randomness. However, it is interesting to see that some structural properties are still preserved, for instance, the region within the marked rectangle. **Bottom row:** We annotate the pair of segments corresponding to the region of interest we discussed above. These segments seem to have contact in 3D space even for the sequence generated with high temperature (`gen2`).

**Figure 5.**
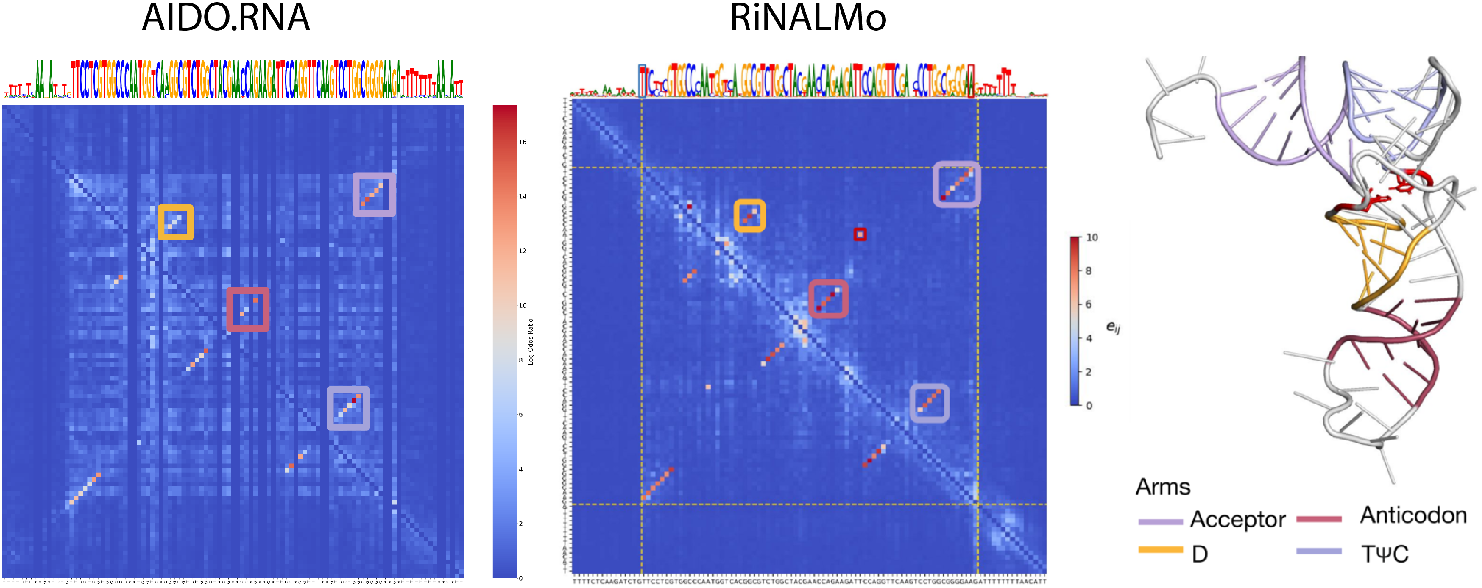
Comparing dependency mapping [74] with AIDO.RNA and RiNALMo for identifying tRNA secondary structure.

### 3.5 High-quality small data is better than low-quality large data for pre-training

From the literature (as shown in Table 22), we find that there is no consensus regarding the ideal dataset for pre-training a versatile RNA FM that can benefit diverse RNA downstream tasks. Uni- RNA [9] employs 1B potential RNA sequences for pre-training, while RiNALMo [10] utilizes 36M ncRNA sequences. The extensive number of sequences used in Uni-RNA’s pre-training is enticing for training a general-purpose RNA foundation model. Although Uni-RNA does not publicly release their pre-training data, the MARS dataset [60] contains similar data sources. We analyze the data and find that approximately 85% of the sequences within MARS are whole-genome shotgun sequences, indicating a significant portion of the data consists of DNA fragments. Consequently, despite the dataset’s substantial size, its quality is considerably low. To explore the effect of noisy data, we pre-train a 1B model utilizing this extensive yet low-quality dataset and compare the model with models trained using a smaller but high-quality dataset. For details of our 1B model’s pre-training data and setting, see Appendix Section F. We compare AIDO.RNA-1B with Uni-RNA, RiNALMo, and our AIDO.RNA-1.6B model on RNA secondary structure prediction, mean ribosome load prediction, and cross-species splice site prediction tasks. As shown in Table 11, our AIDO.RNA-1B trained on a low-quality large dataset performs worse than RiNALMo trained on a high-quality small dataset. This suggests that a large pre-training dataset does not necessarily benefit downstream tasks if the data quality is low and the data distribution differs from the specific task. Moreover, we observe that AIDO.RNA-1B achieves SOTA performance on cross-species splice site prediction, outperforming both RiNALMo and AIDO.RNA-1.6B, which were trained on the RNAcentral database. This improvement may be attributed to the similarity between the input sequences for this task and the majority of sequences in MARS50. Aligning the pre-training dataset with the downstream task dataset holds promise for enhancing downstream task performance.

**Table 11:** Downstream task performance comparison regarding the pre-training dataset. The result of RiNALMo on cross-species splice site prediction is reproduced by us.

| Model | Pre-training data | SS TS0 (F1) | MRL($R^2$) | CSP (F1) |
| --- | --- | --- | --- | --- |
| Uni-RNA [9] | 1B noisy RNA sequences/DNA fragments | - | 0.85 | 0.947 |
| AIDO.RNA-1B(ours) | 886M noisy RNA sequences/DNA fragments | 0.69 | 0.86 | **0.967** |
| RiNALMo [10] | 36M high-quality ncRNA sequences | 0.75 | **0.86** | 0.945 |
| AIDO.RNA-1.6B(ours) | 42M high-quality ncRNA sequences | **0.79** | **0.86** | 0.949 |

**Table 12:** RNA type distribution of training, validation, and test set.

| RNA type | # train | # valid | # test | train ratio | valid ratio | test ratio |
| --- | --- | --- | --- | --- | --- | --- |
| rRNA | 28,913,600 | 10,000 | 10,000 | 69.7% | 12.5% | 12.5% |
| tRNA | 3,543,484 | 10,000 | 10,000 | 8.5% | 12.5% | 12.5% |
| lncRNA | 3,531,218 | 10,000 | 10,000 | 8.5% | 12.5% | 12.5% |
| misc_RNA | 2,463,079 | 10,000 | 10,000 | 5.9% | 12.5% | 12.5% |
| sRNA | 468,084 | 5,000 | 5,000 | 1.1% | 6.3% | 6.3% |
| pre_miRNA | 360,711 | 5,000 | 5,000 | 0.9% | 6.3% | 6.3% |
| ncRNA | 348,184 | 5,000 | 5,000 | 0.8% | 6.3% | 6.3% |
| snRNA | 302,658 | 5,000 | 5,000 | 0.7% | 6.3% | 6.3% |
| snoRNA | 281,057 | 5,000 | 5,000 | 0.7% | 6.3% | 6.3% |
| piRNA | 209,806 | 5,000 | 5,000 | 0.5% | 6.3% | 6.3% |
| SRP_RNA | 200,854 | 5,000 | 5,000 | 0.5% | 6.3% | 6.3% |
| others | 879,659 | 5,000 | 5,000 | 2.1% | 6.3% | 6.3% |
| total | 41,502,394 | 80,000 | 80,000 | 100.0% | 100.0% | 100.0% |

**Table 13:**
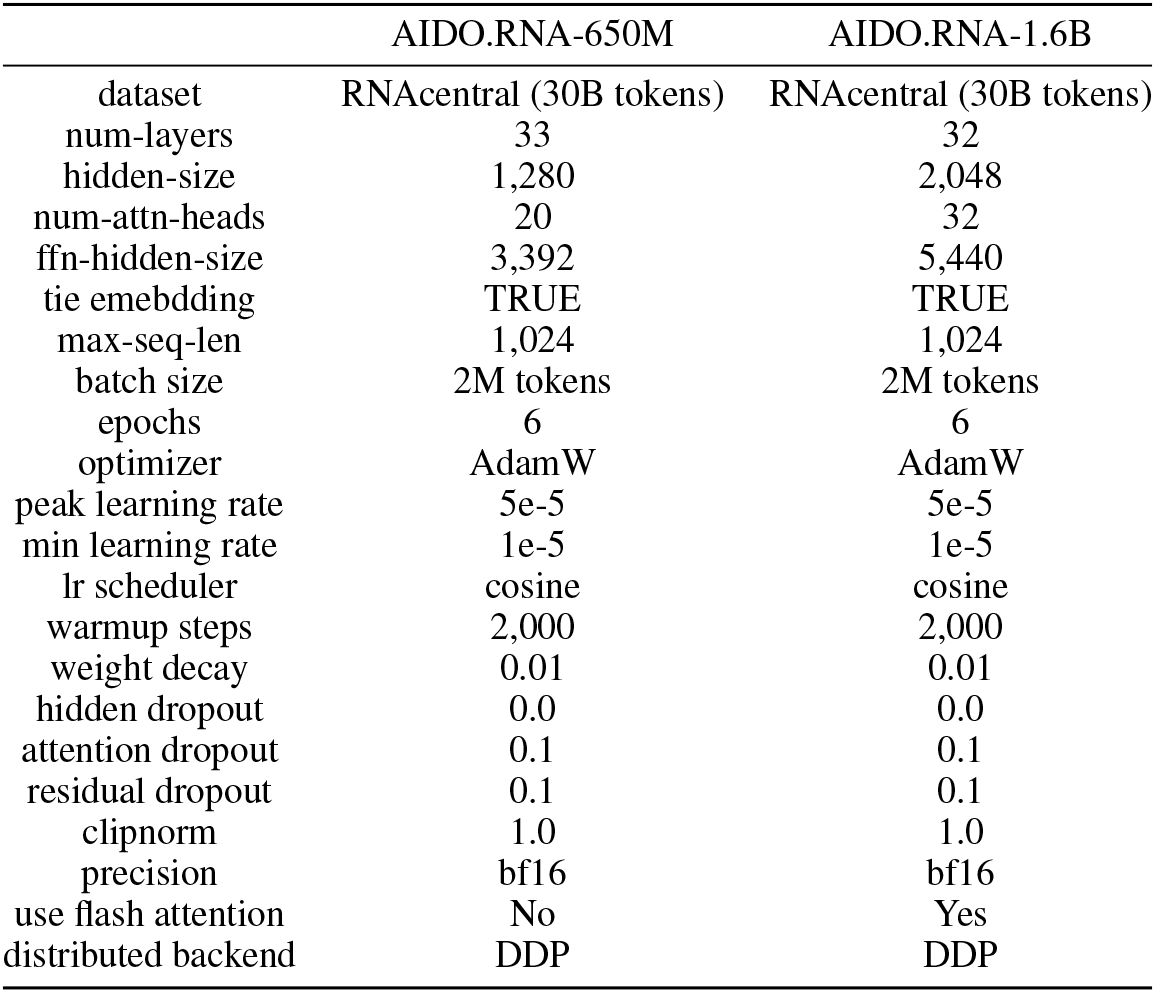
Hyperparameters for training AIDO.RNA models.

## 4. Conclusions and future work

In this work, we present AIDO.RNA, the largest general-purpose RNA foundation model to-date and a key module in an AI-driven Digital Organism. AIDO.RNA excels in a diverse set of RNA under- standing and generation tasks. We find that high-quality data is crucial for pre-training powerful RNA foundation models. We also find that domain-adaptive pre-training yields significant performance gain in the target domain, further emphasizing the importance of pre-training data. By collecting more high-quality RNA sequences and better mixing different domains, it is promising to build a larger and stronger RNA foundation model. We leave this part for future work.

## Appendix

### A. Pre-training data and hyperparameters

We pre-train two RNA foundation models with different model size using ncRNA sequences from RNAcentral database. Table 12 shows the data distribution of our pre-training data. Table 13 shows the key configurations of our models.

### B RNA sequence understanding tasks

We build a comprehensive benchmark to fully evaluate our RNA foundation model by integrating tasks from the literature [9, 7, 8]. As shown in Table 14, the benchmark encompasses a total of 26 subtasks from 9 distinct tasks, including RNA structure, function prediction, and mRNA-related prediction tasks critical for mRNA therapeutic design.

**Table 14:** Overview of RNA sequence understanding tasks. Bold denotes the most important task at the RNA level.

| Category | Task | #subtask | Input | Task formulation |
| --- | --- | --- | --- | --- |
| Structure | **RNA secondary structure prediction** | 2 | ncRNA | pairwise token-level binary classification |
| mRNA-related | **Translation efficiency prediction** | 3 | 5' UTR | seq-level regression |
|  | Expression level prediction | 3 | 5' UTR | seq-level regression |
|  | Mean ribosome load prediction | 1 | 5' UTR | seq-level regression |
|  | Transcript abundance prediction | 7 | CDS | seq-level regression |
|  | Protein abundance prediction | 5 | CDS | seq-level regression |
| Function | Cross-species splice site prediction | 2 | pre-mRNA | seq-level binary classification |
|  | ncRNA family classification | 2 | ncRNA | seq-level multi-class classification |
|  | RNA modification site prediction | 1 | RNA | seq-level multi-label classification |

### B.1 Task datasets

#### RNA secondary structure prediction

In this study, we employed two benchmark datasets for secondary structure prediction, as developed by Singh et al. (2019)[31] and Szikszai et al. (2022)[32]. The former dataset was derived from bpRNA-1m [33] by filtering out sequences longer than 500 bases and applying an 80% sequence similarity cut-off. This preprocessed dataset was divided into three splits: TR0 for training with 10,814 samples, VL0 for validation with 1,300 samples, and TS0 for testing with 1,305 samples.

For generalization assessment, we used the dataset by Szikszai et al. (2022)[32], consisting of 3,865 RNAs from nine families. This dataset was generated from the Archive-II dataset[61] by filtering out sequences longer than 512 nucleotides. It was then split into nine subsets, each time leaving out one family for evaluation while using the remaining families for training and validation. In Table 15, we show the statistics of each family in filtered version of ArchiveII dataset.

**Table 15:** Family-wise statistics of RNA sequences in filtered ArchiveII.

| RNA family | Mean length | Total count |
| --- | --- | --- |
| 5S rRNA | 119 | 1283 |
| SRP RNA | 180 | 918 |
| tRNA | 77 | 557 |
| tmRNA | 366 | 462 |
| RNase P RNA | 332 | 454 |
| Group I Intron | 375 | 74 |
| 16S rRNAa | 317 | 67 |
| Telomerase RNA | 438 | 35 |
| 23S rRNAa | 326 | 15 |
| Mean | 281 | 429 |
| Total | - | 3865 |

#### Translation efficiency and expression level prediction

We use the data from UTR-LM [7], which contains three datasets gathered from human muscle tissue (Muscle), human prostate cancer cell line PC3 (PC3), and human embryonic kidney 293T cell line (HEK) ^3^. The Muscle, PC3, and HEK datasets contain 1,257, 12,579, and 14,410 samples correspondingly. Each sample in these datasets includes a UTR sequence, a translation efficiency label, and an expression level label. The expression level label is measured using RNA-sequencing RPKM, where RPKM refers to reads per kilobase of transcript per million mapped reads. The translation efficiency label is measured by dividing the Ribo-seq RPKM by the RNA-sequencing RPKM. Note that in the datasets released by UTR-LM, all labels are transformed into the natural logarithm space. Following UTR-LM, we use these datasets for training and testing via 10-fold cross-validation. Table 16 summarizes the data statistics.

**Table 16:** Statistics of translation efficiency and expression level prediction datasets. “TE” denotes translation efficiency, “EL” denotes expression level.

| Cell line | Samples | Seq len |  |  | TE label |  | EL label |  |
| --- | --- | --- | --- | --- | --- | --- | --- | --- |
|  |  | Min | Max | Mean | Mean | Std | Mean | Std |
| Muscle | 1,257 | 45 | 100 | 91 | -0.26 | 1.46 | 3.12 | 1.30 |
| PC3 | 12,579 | 45 | 100 | 91 | -0.36 | 1.01 | 3.07 | 1.09 |
| HEK | 14,410 | 45 | 100 | 91 | -0.65 | 1.08 | 3.01 | 1.01 |

#### Mean ribosome load prediction

For this task, we use the same benchmark dataset used by previous best-performing methods, which is a large-scale synthetic Human 5’UTR library [40] consisting of 83,919 5’UTRs (untranslated regions) spanning 75 distinct lengths, each paired with its associated MRLs. To ensure balanced validation, 7,600 sequences are uniformly sampled at each length (namely Random7600), while the remaining data is allocated for training purposes. Note that we use the same splitting as the previous studies for a fair comparison [12, 40]. Furthermore, an extra dataset containing 7,600 authentic human 5’UTRs, distributed in a similar manner as the synthetic collection, is employed as the test set (namely Human7600).

#### Transcript abundance prediction

We use the public data from caLM [8] for transcript abundance prediction ^4^. It contains seven datasets from *A. thaliana, D. melanogaster, E*.*coli, H. sapiens, S. cerevisiae, H. volcanii* and *P. pastoris*, respectively. The abundance label is the natural logarithm of the transcript count per million, ranging between -16 and 16. Table 17 shows the overall data statistics for each organism.

**Table 17:** Statistics of transcript abundance prediction datasets.

| Organism | Sample | Seq len |  |  | Label |  |  |
| --- | --- | --- | --- | --- | --- | --- | --- |
|  |  | Min | Max | Mean | Min | Max | Mean |
| *A. thaliana* | 12,664 | 120 | 16182 | 1312 | -3.32 | 16.13 | 2.55 |
| *D. melanogaster* | 9,836 | 132 | 45318 | 1718 | -3.32 | 13.59 | 3.42 |
| *E. coli* | 3,528 | 117 | 7077 | 971 | -5.81 | 14.16 | 4.49 |
| *H. sapiens* | 5484 | 117 | 16941 | 1578 | 1.85 | 13.63 | 5.21 |
| *S. cerevisiae* | 5,448 | 105 | 14733 | 1408 | -3.32 | 14.30 | 5.72 |
| *H. volcanii* | 3,189 | 135 | 6717 | 892 | -6.04 | 13.99 | 5.17 |
| *P. pastoris* | 4,682 | 207 | 14811 | 1494 | -16.30 | 15.44 | 5.11 |

#### Protein abundance prediction

We use the public data from caLM [8] for protein abundance prediction ^5^. It contains five datasets from *A. thaliana, D. melanogaster, E*.*coli, H. sapiens*, and *S. cerevisiae*, respectively. The abundance labels are the estimated number of copies per cell annotated in PAXdb [45], ranging between 0 and 10^5^. Table 18 shows the overall data statistics for each organism.

**Table 18:**
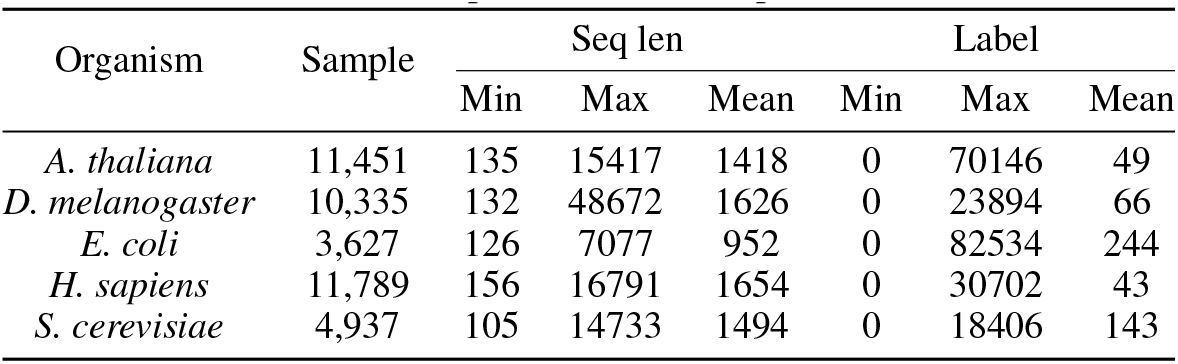
Statistics of protein abundance prediction datasets.

#### Cross-species splice site prediction

We use the data from Spliceator, which contains confirmed error-free splice sites from more than 100 eukaryotic species [46]. In specific, we use the acceptor and donor datasets from GS-1 subset ^6^ for training and validation. For testing, we use data from four different species that are not shown in the training set, including zebrafish, fruit fly, worm, and plant _7_. Each sample in the dataset is a 400nt sequence centered on a splice/non-splice site. Table 19 shows the overall data statistics.

**Table 19:** Statistics of splice site prediction datasets.

|  | Train | Valid | Test |  |  |  |
| --- | --- | --- | --- | --- | --- | --- |
|  |  |  | Zebrafish | Fly | Worm | Plant |
| Acceptor | 17,723 | 4,431 | 20,000 | 20,000 | 20,000 | 20,000 |
| Donor | 17,556 | 4,389 | 20,000 | 20,000 | 20,000 | 20,000 |

#### Non-coding RNA family classification

We use the preprocessed Rfam-novel dataset ^8^ from [48], which contains 105,864/17,324/25,342 in the train/valid/test sets correspondingly. The dataset contains 88 Rfam classes, with an imbalanced class distribution in the training set. The ncRNA sequence length with 0 boundary noise is ≤ 200. When the boundary noise increases to ≤ 200%, the sequence length increases two times.

#### RNA modification site prediction

We use the processed data from MultiRM [50], which is obtained from 20 epi-transcriptome profiles generated from 15 different base-resolution technologies for 12 different types of RNA modifications. The 12 modifications are *Am, Cm, Gm, Tm, m*^1^*A, m*^5^*C, m*^5^*U* , *m*^6^*A, m*^6^*Am, m*^7^*G*, Φ, and *I*. The train/valid/test data contains 304,661/3,599/1,200 samples respectively, with a sequence length of 1,001. Negative sites were chosen at random from the unmodified bases within the same transcript that also contains the positive sites. Table 20 shows label distribution for each modification type.

**Table 20:** Number of positive samples for each modification site in the RNA modification site prediction dataset.

| | *Am* | *Cm* | *Gm* | *Tm* | $m^1A$ | $m^5C$ | $m^5U$ | $m^6A$ | $m^6Am$ | $m^7G$ | $\Phi$ | *I* |
| --- | --- | --- | --- | --- | --- | --- | --- | --- | --- | --- | --- | --- |
| Train | 1,391 | 1,678 | 1,271 | 2,053 | 16,146 | 3,007 | 3,496 | 64,978 | 2,247 | 836 | 2,937 | 52,418 |
| Valid | 150 | 150 | 150 | 150 | 150 | 150 | 150 | 150 | 150 | 150 | 150 | 150 |
| Test | 50 | 50 | 50 | 50 | 50 | 50 | 50 | 50 | 50 | 50 | 50 | 50 |

**Table 21:** Inverse folding performance comparison on 14 RNA structures of interest identified by Das *et al*. [59]. All of these 14 RNAs belong to the test set. The results by gRNAde [23] were produced by the model checkpoint provided in the official github repository (source: https://github.com/chaitjo/geometric-rna-design, checkpoint file name gRNAde_ARv1_1state_all.h5. Last accessed: Sept 25, 2024).

| PDB ID | Description | gRNAde | gRNAde +<br>AIDO.RNA-cDiffusion |
| --- | --- | --- | --- |
| 1CSL | RRE high affinity site | 50.0 | 53.846 |
| 1ET4 | Vitamin B12 binding RNA aptamer | 42.857 | 45.714 |
| 1F27 | Biotin-binding RNA pseudoknot | 50.0 | 44.444 |
| 1L2X | Viral RNA pseudoknot | 59.259 | 62.963 |
| 1LNT | RNA internal loop of SRP | 65.0 | 60.0 |
| 1Q9A | Sarcin/ricin domain from E.coli 23S rRNA | 88.889 | 92.593 |
| 4FE5 | Guanine riboswitch aptamer | 41.791 | 46.269 |
| 1X9C | All-RNA hairpin ribozyme | 40.0 | 48.333 |
| 1XPE | HIV-1 B RNA dimerization initiation site | 60.87 | 65.217 |
| 2GCS | Pre-cleavage state of glmS ribozyme | 45.902 | 47.541 |
| 2GDI | Thiamine pyrophosphate-specific riboswitch | 58.974 | 60.256 |
| 20EU | Junctionless hairpin ribozyme | 38.095 | 33.333 |
| 2R8S | Tetrahymena ribozyme P4-P6 domain | 70.886 | 68.987 |
| 354D | Loop E from E. coli 5S rRNA | 60.0 | 65.0 |
| Average |  | 55.18 | 56.74 |

**Table 22:**
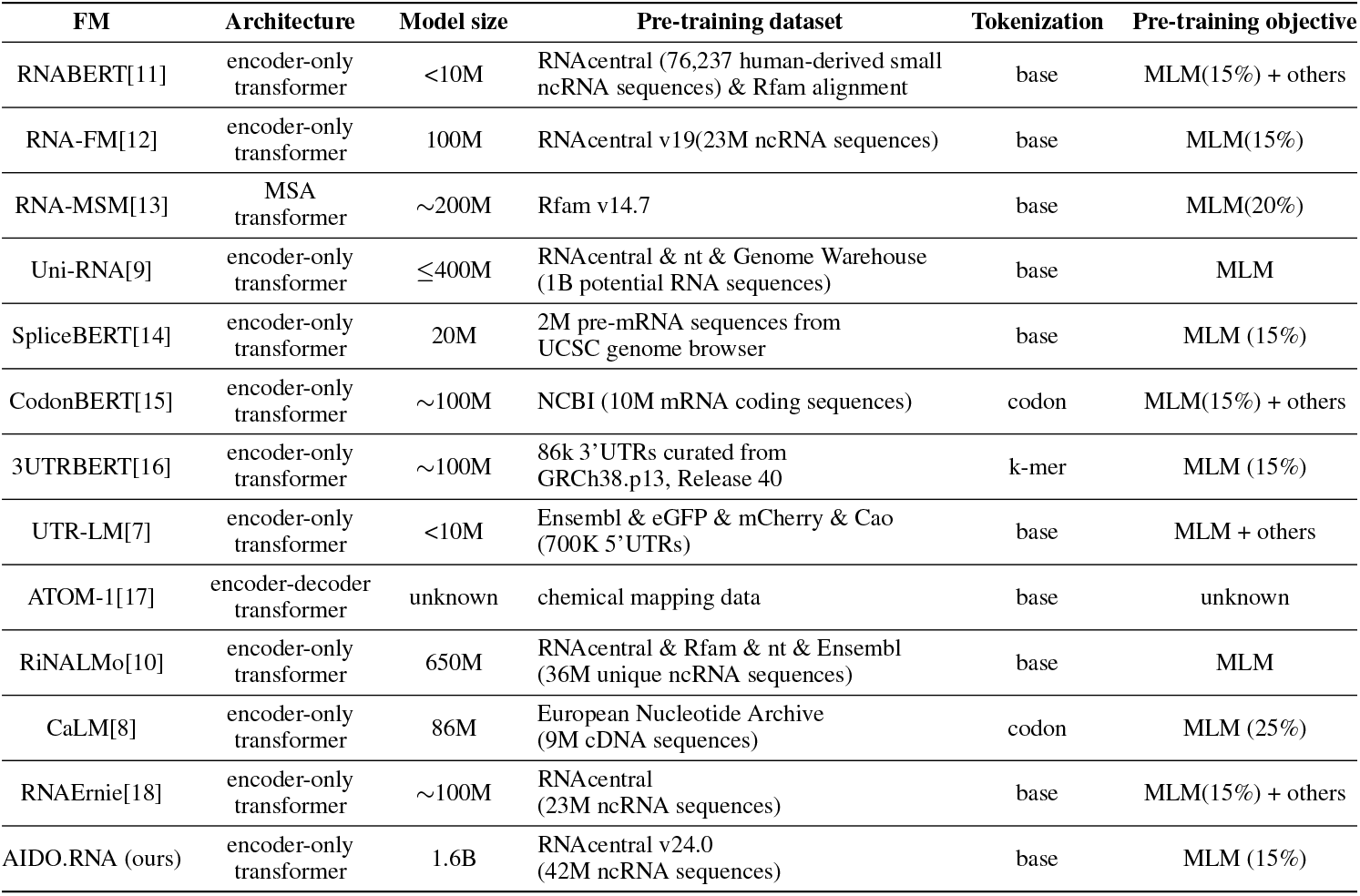
Related work of RNA foundation models.

### B.2 Fine-tuning settings

#### RNA secondary structure prediction

We generate pair representations by applying outer concatenation to the language model’s outputs, concatenating the representation of nucleotide *j* with that of nucleotide *i* for each pair (*i, j*). In the prediction head the concatenated representation is first linearly projected to an embedding dimension of 64. This is followed by two bottlenecked ResNet-2D blocks and a convolution layer, all with 64 kernels of size 3 and followed by instance normalization and ReLU activation. The output matrix represents pairing probability logits for nucleotide pairs, with binary cross-entropy loss calculated only for elements above the main diagonal due to the symmetry of secondary structures. To train the model, we use AdamW optimizer with weight decay 0.01. To prevent the model from overfitting, we regularize both the language model and the prediction head with dropout rate of 0.1. Fine-tuning involved a gradual unfreezing method, starting with training the prediction head for the first three epochs, then unfreezing three additional layers every three epochs over a total of 60 epochs with a batch size of 4, and a learning rate decreasing from 10^*−*4^ to 10^*−*5^. A greedy algorithm was used to convert base pairing probabilities into secondary structures, prioritizing the highest probabilities and excluding conflicting pairs, avoiding non-canonical pairings and sharp hairpin loops (|*i*−*j*|*<* 4). The classification threshold was tuned on the validation set to balance the pairing ratio.

#### Translation efficiency and expression level prediction

Since translation efficiency and expression level prediction are sequence-level regression tasks, we take the mean pooling of the output of the transformer model as sequence representation and add a two-layer MLP with a hidden dimension of 512 as the prediction head. Mean square error (MSE) is then used as the loss for both tasks. We fully fine-tune the model using AdamW with a peak learning rate of 1e-5 and weight decay of 0.01. The dropout probability is set to 0.1. For the Muscle dataset, the batch size is set to 8 while for the PC3 and HEK datasets, the batch size is set to 32. We train the model for 30 epochs and select the best checkpoint based on the validation score. To make results comparable with UTR-LM[7], we use 10-fold cross-validation for each cell line on both translation efficiency prediction and expression level prediction tasks.

#### Mean ribosome load prediction

Predicting mean ribosome load is also a regression task performed at the sequence level. The prediction head comprises a linear projection into 256 dimensions and nine ResNet-1D blocks. Each block includes two convolution layers with 256 kernels of size 3, followed by instance normalization and an ELU activation function. The model is also regularized with dropout at rate 0.1. The MRL targets were standardized using the mean and standard deviation of the training MRL values. The model underwent fine-tuning for 60 epochs. During the first 3 epochs, only the prediction head was trained. The learning rate, similar to that used in RNA secondary structure prediction, started at 10^*−*4^ and linearly decreased to 10^*−*5^ over the first 5000 training steps before remaining constant. The batch size for the training process was set to 64. We train the model with AdamW optimizer for this task as well.

#### Protein abundance and transcript abundance prediction

To adapt our model to the CDS domain, we continue to pre-train our model on 9 million CDS sequences released by CaLM [8]. We trained our model for 13,000 steps with a peak learning rate of 5e-5 and a mask ratio 0.25. For both tasks, we take the mean pooling of the transformer’s output as sequence representation and add a two-layer MLP with a hidden dimension of 512 as the prediction head. Mean square error (MSE) is used as the loss. Following CaLM, we perform 5-fold cross-validation on each organism dataset. We use LoRA fine-tuning with rank=32 and alpha=64. The trainable parameters amount to 9 million, which constitutes 0.58% of the total size of the pre-trained model. We use AdamW with a weight decay of 0.01. Both the hidden layer and LoRA dropout probabilities are set to 0.1. The batch size is set to 16. We train the model for 15 epochs and select the best checkpoint based on the validation score. For the protein abundance prediction task, we convert the label *y* to natural logarithm space by using *log*(1 + *y*). The peak learning rate is set to 3e-4 for both tasks. Since the sequences on each organism are generally long, we truncate each sequence to 1,024 nucleotides as input.

#### Cross-species splice site prediction

For this sequence-level binary classification task, we take the `[CLS]` embedding from the output of the transformer as sequence representation and add a three-layer MLP with hidden dimensions of 512 and 128 as the prediction head. Cross-entropy loss is used as the loss. We use LoRA fine-tuning with rank=32 and alpha=64. The trainable parameters amount to 9.6 million, which constitutes 0.59% of the total size of the pre-trained model. We train the model using AdamW with a peak learning rate of 2.5e-4 and weight decay of 0.01. The batch size is set to 32. Both the hidden layer and LoRA dropout probabilities are set to 0.1. We train the model for 10 epochs and select the best checkpoint based on the validation score.

#### Non-coding RNA family classification

For this sequence-level classification task, we take the mean pooling of the output of the transformer as sequence representation and add a two-layer MLP with a hidden dimension of 512 as the prediction head. Cross-entropy loss is used as the loss. We use LoRA fine-tuning with rank=16 and alpha=32. The trainable parameters amount to 5 million, which constitutes 0.33% of the total size of the pre-trained model. We train the model using AdamW with a peak learning rate of 4e-4 and weight decay of 0.01. The batch size is set to 64. Both the hidden layer and LoRA dropout probabilities are set to 0.1. We train the model for 15 epochs and select the best checkpoint based on the validation score.

#### RNA modification site prediction

For this sequence-level multi-label classification task, we take the `[CLS]` embedding from the output of the transformer as sequence representation and add a two-layer MLP with a hidden dimension of 512 as the prediction head. Binary cross-entropy loss is computed for each modification site at the same time. We use LoRA fine-tuning with rank=16 and alpha=32. We train the model using AdamW with a peak learning rate of 4e-4 and weight decay of 0.01. The batch size is set to 64. Both the hidden layer and LoRA dropout probabilities are set to 0.1. We train the model for 10 epochs and select the best checkpoint based on the validation score.

### C RNA inverse folding

RNA inverse folding, also referred to as structure-conditioned RNA design, aims to generate RNA sequences that fold into a predefined 3D structure [23, 58]. It is an inverse task of RNA structure prediction which aims to predict the structure based on a given sequence [62]. In RNA inverse folding, the challenge lies in identifying sequences that can reliably adopt the desired structure [63, 51]. In particular, we focus on designing sequences with known RNA backbone structure [23, 63]. This task is vital for synthetic biology and nanotechnology applications [64], where specific RNA structures are needed to perform essential biological functions, such as acting as molecular switches [65], facilitating biochemical interactions [66], or serving as scaffolds for molecular assemblies [67].

The complexity of RNA inverse folding stems from the intricate relationship between RNA sequence and structure. RNA molecules exhibit a high degree of structural variability, and even small changes in sequence can lead to significantly different folding patterns [63]. Therefore, the challenge for computational models is to identify sequences that thermodynamically favor the target structure while avoiding undesired or alternative configurations. This task requires advanced algorithms capable of optimizing sequences based on both stability and specificity. Recent advances in computational modeling, particularly those leveraging deep generative models, have significantly improved the accessibility and effectiveness of RNA inverse folding approaches [23, 63, 51].

#### C.1 Method

Building on a state-of-the-art method gRNAde [23], we demonstrate how AIDO.RNA can be used to enhance RNA inverse folding performance. We conduct two separate experiments to showcase the capabilities of AIDO.RNA: one focusing on adaptation with conditional diffusion and the other on zero-shot generation. In the diffusion-adapted model, AIDO.RNA is fine-tuned for the inverse folding task, while in the zero-shot setting, we evaluate its refinement capabilities without any adaptation.

##### C.1.1 Masked diffusion for RNA sequence generation

We aim to approximate a data distribution *q*(*x*) by training a diffusion model, by first iteratively adding noise to a sample *x*_0_∼*q*(*x*) for *T* discrete steps (forward process) that results in a sample with entire noise *x*_*T*_ ; and then training a model, parameterized by *θ*, to denoise *x*_*T*_ iteratively to retrieve the original signal *x*_0_ (reverse process). For continuous signals, such as image or audio, at any time step *t* ∈ [0, *T* ], the sample *x*_*t*_ can be assumed as a linear combination of the original signal *x*_0_ and Gaussian noise *ϵ* ∼ 𝒩 (0, 1) as follows:

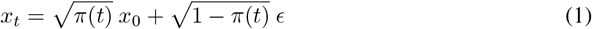

where *π*(*t*) ∈ [0, 1] is a monotonically decreasing function of *t* [68]. The model learns a marginal distribution *p*_*θ*_(*x*_*t*−1_ |*x*_*t*_), which aims to approximate the true transition probability *q*(*x*_*t*−1_|*x*_*t*_, *x*_0_) of estimating a less noisy variant *x*_*t*−1_ given a relatively more noisy variant *x*_*t*_. Assume we have *x*_*t*_ = *x*_0_, *π*(*t*) = 1 at *t* = 0, and *x*_*t*_ = *x*_*T*_ = *ϵ* ∼ 𝒩 (0, 1), *π*(*t*) = 0 at *t* = *T* . For discrete signals like RNA sequence, however, it is infeasible to represent *x*_*T*_ as a sample from standard normal distribution. To address this issue, we represent *x*_*T*_ by *absorbing state* [69, 56] that contain no data-specific signal, analogous to pure Gaussian noise. Following [69], we use the `[MASK]` token as the absorbing state.

##### Training objective

We adopt the formulation proposed by [56] as our masked diffusion model training objective. It is a negative evidence lower bound on log likelihood (NELBO) [69] and can be decoupled into three disjoint objectives for reconstruction 𝒥_*recon*_, diffusion 𝒥_*diff*_ , and prior 𝒥_*prior*_.

As derived by [69, 56, 57], for diffusion directly on data samples *x*, it is possible to show that 𝒥_*recon*_ = 0, 𝒥_*prior*_ = 0. Given this, NELBO for discrete times step *T* can be simplified as follows:

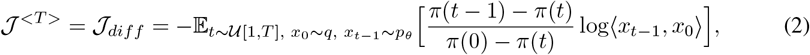

where 𝒰 [1, *T* ] is a uniform distribution integers between 1 and *T* , and ⟨*x*_*t−*1_, *x*_0_⟩ computes the similarity between *x*_*t−*1_ and *x*_0_. In our experiment, we use cross-entropy loss between *x*_0_ and *x*_*t−*1_, ℒ_*CE*_(*x*_0_, *x*_*t−*1_), for − log⟨*x*_*t−*1_, *x*_0_⟩. As shown in [70], we can get a tighter bound on 𝒥 _*<T>*_ with higher number of diffusion steps T. When *T* ⟶ ∞, Equation 2 becomes follows:

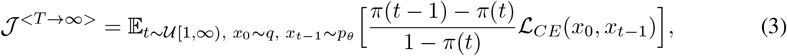

where *π*(0) = 1. Note that for *T* ⟶ ∞, *π*(*t* − 1) ⟶ *π*(*t*), i.e., the change in *π*(*t*) at any time *t* should be infinitesimal. Besides, we have *π*(*t* − 1) − *π*(*t*) *>* 0 since *π*(*t*) is monotonically decreasing. With *T* ⟶ ∞, we can represent this change with the negative time-derivative of *π*(*t*) at time *t*,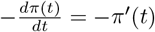. This leads to the continuous-time likelihood bound as follows:

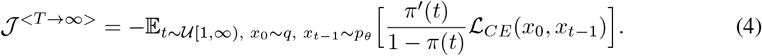

As shown by [56], the choice of *π*(*t*) has insignificant effect on the overall performance of the training algorithm. We adopt 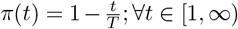 as our noise schedule for its simplicity and efficiency. This further simplifies Equation 4 as follows:

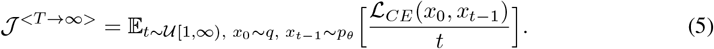

##### Intuition behind the objective function

Note that the loss computed on any sample *x*_*t*_ is now inversely proportional to *t*. Intuitively, if *t* is large, *x*_*t*_ is more noisy and hence it can potentially lead to many varieties of reconstructed samples 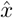 from *q*(*x*), i.e., all of them can be valid. However, with ℒ_*CE*_(*x*_0_, *x*_*t*_) loss we are always pushing the *x*_*t*_ to be more similar to *x*_0_, i.e., encouraging less diversity in generation, which is only expected if *x*_*t*_ is already very similar to *x*_0_ (when *t* is smaller). To address this conflict, the loss ℒ_*CE*_(*x*_0_, *x*_*t*_) is down-weighted by the factor 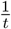 .

##### C.1.2 Adaptation with conditional diffusion

For a given 3D RNA structure, we begin by taking the predicted sequence and structural embedding by gRNAde, *S*_0_ and *e*^*st*^, respectively. We then mask out 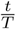 (where *t* ∼ 𝒰 [1, *T* ]) portion of *S*_0_, that produces *x*_*t*_, which can be assumed as a noisy variant of the expected RNA sequence. We then pass *x*_*t*_ through AIDO.RNA’s encoder that produces sequence embedding 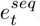. Then *e*^*st*^ and 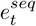 are processed by an adaptor module [71, 72], which in our design is a multi-head self-attention layer [19] with bottleneck [73], that generates a new estimate of the RNA sequence *x*_*t* − 1_. Note that here the AIDO.RNA and the adaptor combined work as the estimated transition function *p*_*θ*_(*x*_*t*−1_ |*x*_*t*_) we discussed in the previous section. During training, we optimize the diffusion objective in Equation 5, where *x*_0_ is the ground truth sequence. After training, we can generate sequences starting from the initial estimate *S*_0_ and structural embedding *e*^*str*^ provided by gRNAde, and applying masking to the top *M* least confident tokens in *S*_0_ (based on predicted class probabilites of the tokens), resulting in a masked sequence *x*_*t*_. We then iteratively denoise this sequence over several steps to obtain 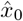 , our final estimate.

##### C.1.3 Zero-shot generation

This method is utilizes the generation approach in the conditional diffusion discussed above; however, it operates entirely in a zero-shot manner. This means we utilize our pre-trained AIDO.RNA without any fine-tuning or the implementation of an adaptor module. Specifically, we begin with the initial estimate *S*_0_ generated by gRNAde. Subsequently, we apply a masking technique to the top *M* least confident tokens in *S*_0_, resulting in a masked sequence *x*_*t*_. Following this, we iteratively denoise the sequence over several steps, gradually refining it until we arrive at 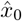 , which serves as our generated sequence. Please note that, unlike the diffusion-adaption settings, here we utilize the full AIDO.RNA architecture (i.e., both the AIDO.RNA’s encoder and its masked language modeling prediction head). This approach ensures that we can leverage the pre-trained capabilities of AIDO.RNA effectively while maintaining simplicity in the process.

#### C.2 Results

### D AIDO.RNA captures structure information through self-supervised pre-training

Self-supervised pre-training is a powerful tool to infer conditional dependencies, which can be probed and cataloged through *in silico* mutagenesis. To assess conditional dependencies, we implement the dependency mapping strategy in [74],

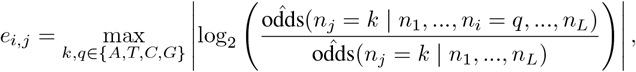

where *e*_*i*,*j*_ are the “pixel” values of the dependency map, *k* and *q* are the key and query nucleotides, *n* is a length *L* DNA sequence, and odds are the odds inferred by the pre-trained RNA Foundation.

While *in silico* mutagenesis studies normally require *O*(*L*^2^) inferences using a supervised model such as Enformer [75] to compute all pairwise interactions, self-supervised models infer the probability of all key mutations under a given query mutation at once, allowing us to compute this dependency mapping with only *O*(*L*) inferences.

### E Related work

Table 22 provides an overview of the literature on pre-training RNA foundation models. These models predominantly use an encoder-only transformer architecture and employ masking language modeling (MLM) as the pre-training objective. The key distinctions among these models are the pre-training dataset and model size. In comparison, our AIDO.RNA is the largest RNA foundation model up-to-date and it is pre-trained on ncRNA sequences from RNAcentral.

### F. Pre-training AIDO.RNA-1B on a vast number of noisy RNA sequences

#### MARS50

To pre-train the 1B model, we use genomic sequences from the MARS database, which contains 1.7 billion nucleotide sequences from various biological databases [60]. The sequences are aligned to a standardized DNA alphabet and undergo filtering and clustering steps. Extremely long and short sequences (exceeding 4,096 bases or below 10 bases) are excluded. We perform a two-step clustering approach using the MMseqs2 algorithm [25]. The first step clusters sequences with at least 90% identity and 80% overlap with the longest sequence in each cluster, resulting in the MARS90 dataset. In the second step, sequences with at least 50% identity and 80% overlap with the longest sequence are clustered, yielding the refined MARS50 dataset. MARS50 comprises 886 million sequences, totaling 344 billion bases, with an average length of 389 bases. Finally, the dataset is randomly split, with 0.2% allocated for validation and another 0.2% for testing purposes.

#### Pre-training

We pre-train the AIDO.RNA-1B model, which consists of 1 billion parameters, using the MARS50 dataset. This model has 36 layers, 32 attention heads, and a hidden size of 1,536. The other architecture hyperparameters are the same as our 1.6B model. During pre-training, we train the model for 145k steps, employing MLM loss with a mask ratio of 15%. We use AdamW optimizer with a peak learning rate of 1.5e-4 and a weight decay of 0.02.

### G AIDO.RNA-650M: our reproduction of RiNALMo

Before our work, the largest RNA foundation model available was RiNALMo [10], which consisted of 650 million parameters. It was pre-trained using 36 million ncRNA sequences collected from the RNAcentral database, Rfam, nt, and Ensembl. It achieved SOTA results on RNA secondary structure prediction. Given its impressive performance on downstream tasks, we set out to pre-train a 650M RNA foundation model using almost the same architecture and training setting as RiNALMo before training our 1.6B model. Since RiNALMo does not release its pre-training data, we use the ncRNA sequences from RNAcentral version 24.0 as described in Section 2, which contains all the ncRNA sequences used in RiNALMo’s pre-training in theory. For model comparison, we fine-tune our 650M model and RiNALMo on RNA secondary structure prediction, translation efficiency prediction, and expression level prediction datasets using the same prediction head and training settings. Table 23 shows the results of our AIDO.RNA-650M and RiNALMo. On all 8 datasets from the three tasks, our model achieves similar results as RiNALMo. These results demonstrate that we successfully reproduce RiNALMo, showing the effectiveness of pre-training on a high-quality ncRNA sequence database.

**Table 23:** Comparsion of downstream task performance between AIDO.RNA-650M and RiNALMo [10].

| Task | Dataset | Metric | RiNALMo | AIDO.RNA-650M (ours) |
| --- | --- | --- | --- | --- |
| Secondary structure | bpRNA TSO | F1-score | 0.772 | **0.778** |
|  | Archive-II | F1-score | 0.720 | **0.743** |
| Translation efficiency | Muscle | Spearman CC | **0.711** | 0.697 |
|  | PC3 | Spearman CC | 0.698 | **0.699** |
|  | HEK | Spearman CC | 0.661 | **0.664** |
| Expression level | Muscle | Spearman CC | **0.698** | 0.688 |
|  | PC3 | Spearman CC | 0.672 | **0.683** |
|  | HEK | Spearman CC | 0.697 | **0.707** |

### H Data and code availability

We have developed the ModelGenerator package to reproduce, apply, and extend the results in this manuscript https://github.com/genbio-ai/ModelGenerator. Pre-trained models and down- stream datasets are also available on Hugging Face at https://huggingface.co/genbio-ai.

### Footnotes

3 https://drive.google.com/drive/folders/190oihtrwCxWjtDCK9kJzyhXPKxbr5xoR

4 https://github.com/oxpig/CaLM/tree/main/data/transcript_abundance

5 https://github.com/oxpig/CaLM/tree/main/data/protein_abundance

6 https://www.lbgi.fr/spliceator/?source=download

7 https://git.unistra.fr/nscalzitti/spliceator/-/tree/master/Data/Benchmarks?ref_type=heads

8 https://github.com/bioinformatics-sannio/ncrna-deep/tree/master/datasets/Rfam-novel

## Notes

### Competing Interest Statement

The authors have declared no competing interest.

